# Loss of E-cadherin enhances IGF1-IGF1R pathway activation and sensitizes breast cancers to anti-IGF1R inhibitors

**DOI:** 10.1101/253278

**Authors:** Alison M. Nagle, Kevin M. Levine, Nilgun Tasdemir, Julie A. Scott, Kara Burlbaugh, Justin Kehm, Tiffany A. Katz, David N. Boone, Britta M. Jacobsen, Jennifer M. Atkinson, Steffi Oesterreich, Adrian V. Lee

## Abstract

**Purpose:** Insulin-like growth factor I (IGF1) signaling regulates breast cancer initiation and progression and associated cancer phenotypes. We previously identified E-cadherin *(CDH1)* as a repressor of IGF1 signaling and in this study examined how loss of E-cadherin affects IGF1R signaling and response to anti-IGF1R therapies in breast cancer.

**Experimental Design:** Breast cancer cell lines were used to assess how altered E-cadherin levels regulate IGF1R signaling and response to two anti-IGF1R therapies. *In situ* proximity ligation assay (PLA) was used to define interaction between IGF1R and E-cadherin. TCGA RNA-seq and RPPA data was used to compare IGF1R activation in estrogen receptor positive (ER+) invasive lobular carcinoma (ILC) and invasive ductal carcinoma (IDC) tumors. ER+ ILC cell lines and xenograft tumor explant cultures were used to evaluate efficacy to IGF1R pathway inhibition in combination with endocrine therapy.

**Results:** Diminished functional E-cadherin increased both activation of IGF1R signaling and efficacy to anti-IGF1R therapies. PLA demonstrated a direct endogenous interaction between IGF1R and E-cadherin at points of cell-cell contact. Increased expression of IGF1 ligand and levels of IGF1R phosphorylation were observed in E-cadherin deficient ER+ ILC compared to IDC tumors. IGF1R pathway inhibitors were effective in inhibiting growth in ER+ ILC cell lines and synergized with endocrine therapy and similarly IGF1R inhibition reduced proliferation in ILC tumor explant culture.

**Conclusions:** We provide evidence that loss of E-cadherin hyperactivates the IGF1R pathway and increases sensitivity to IGF1R targeted therapy, thus identifying the IGF1R pathway as a potential novel target in E-cadherin deficient breast cancers.

**STATEMENT OF SIGNIFICANCE:** IGF1R signaling is an attractive therapeutic target in breast cancer due to its regulation of proliferation, migration, and invasion. However, clinical trials targeting IGF1R have largely been unsuccessful due to lack of biomarkers to stratify patients for therapeutic response. In this study, we demonstrate loss of E-cadherin as a potential biomarker for response to anti-IGF1R therapy, and show efficacy of IGF1R inhibition in ER+ ILC in combination with endocrine therapy. Patients with ER+ ILC have poorer long-term outcomes than patients with ER+ IDC and have a propensity for increased late recurrences, highlighting the need for improved therapeutic strategies for this subtype of breast cancer. Here, we credential IGF1R inhibition as a novel therapeutic strategy in combination with endocrine therapy for the treatment of ER+ ILC.

## INTRODUCTION

IGF1 is a circulating endocrine hormone that is a major regulator of organismal growth and development^1^. IGF1, in combination with estrogen, is essential for normal mammary gland development, and this pathway is deregulated in the initiation and progression of breast cancer^2–5^. Many studies, including from our laboratory, have shown the ability of the IGF1 receptor (IGF1R) to promote mammary tumorigenesis and metastasis^6–9^. Additionally, we showed that when constitutively activated, IGF1R transformed mammary epithelial cells, increased migration and invasion, and induced epithelial to mesenchymal transition (EMT) via the NFkB pathway and upregulation of Snail^6,10^.

Based on these observations, both small molecule tyrosine kinase inhibitors and monoclonal antibodies against IGF1R were tested in clinical trials in breast cancer. Unfortunately, although as many as 50% of breast tumors express IGF1R^11^, these trials only identified a small subset of patients showing a therapeutic response to IGF1R targeted therapy, suggesting that predictive biomarkers are required to identify which patients’ tumors will be responsive^12–15^.

We previously developed an IGF1-signature (IGF-sig) based on microarray analyses, and more recently reported a novel computational method to identify putative biomarkers of IGF1 signaling using a systems biology approach^16^. The latter was based on a proteomic screen using reverse phase protein array (RPPA) on 21 breast cancer cell lines stimulated with IGF1 over a time course^17^. This computational model identified E-cadherin as a putative regulator of IGF1 signaling, and data in the present study indicate that loss of E-cadherin expression can directly increase IGF1R pathway activation and associated phenotypes in breast cancer. Insight into how E-cadherin regulates IGF1R is necessary to aid in our understanding of the oncogenic signaling network, specifically because the loss of E-cadherin i) is implicated in the ability of tumor cells to escape the primary tumor to potentially seed metastatic lesions and ii) is transcriptionally repressed and/or genetically lost in subsets of breast tumors^18–22^.

One such subtype of breast cancer with diminished E-cadherin expression is invasive lobular breast carcinoma (ILC), accounting for 10-15% (~30,000 cases/year in the US) of total breast cancer cases. ILC is defined by the loss of functional E-cadherin *(CDH1),* which occurs in 95% of ILC due to truncating mutations, loss of heterozygosity, and transcriptional repression^23,24^. Due to the loss of E-cadherin protein, ILC cells grow in linear patterns throughout the breast tissue, lacking the ability to form adherens junctions, in contrast to the solid mass growth of the most frequent subtype of breast cancer, invasive ductal breast carcinoma (IDC)^25^. Interestingly, one of the most IGF1 responsive cell lines in our above-referenced proteomic data set was a human ILC cell line, MDA-MB-134-IV, that lacks E-cadherin protein expression and cell-cell junctions^17^. In this study, we characterize the regulation of IGF1R by E-cadherin, and provide evidence that inhibition of IGF1R in E-cadherin deficient breast cancers could potentially serve as an effective therapeutic strategy.

## MATERIALS AND METHODS

### Cell culture

Cell lines were authenticated (most recent date listed [ ]) by the University of Arizona Genetics Core and mycoplasma tested (Lonza #LT07-418). Lab stocks were made following authentication and used for this study. MCF-7 (ATCC; DMEM+10% FBS [06/29/16]), T47D (ATCC; RPMI+10% FBS [02/08/17]), ZR75.1 (ATCC; RPMI+10% FBS [10/13/16]), MDA-MB-231 (ATCC; DMEM+10% FBS [10/13/16]), MDA-MB-134-IV (ATCC; 50/50 DMEM/L15+10% FBS [02/08/17]), SUM44PE (Asterand; DMEM/F12+2% CSS with 5ug/ml insulin, 1ug/ml hydrocortisone, 5mM ethanolamine, 5ug/ml transferrin, 10nM triodothyronime, and 50nM sodium selenite [02/08/17 - no reference profile exists in database]), and BCK4^26^ (MEM+5% FBS with 1nM insulin and 1x NEAA [10/13/16 - no reference profile exists in database) cells were cultured with indicated media conditions.

### Transient siRNA transfection

Cells were reverse transfected with 25nM final concentration of siGENOME human SMARTpool control siRNA (Dharmacon #D-001206) or siGENOME human SMARTpool CDH1 siRNA (Dharmacon #M-003877-02) using Lipofectamine RNAiMAX (Invitrogen #13778) protocol for 48 hours. For IGF1 (GroPep BioReagents #CU100) stimulation, cells were serum starved overnight and pulsed with IGF1 (1nM, 10nM, or 100nM) for 10 minutes.

### Stable shRNA infection

Stable CDH1 knockdown T47D cells were generated using a retro-viral infection of Renilla control (shSCR [5’ TGCTGTTGACAGTGAGCGCAGGAATTATAATGCTTATCTATA GTGAAGCCACAGATGTATAGATAAGCATTATAATTCCTATGCCTACTGCCTCGGA]) and two CDH1 (sh-1 [5’ TGCTGTTGACAGTGAGCGCAAGTGTGTTCATTAATGTTTATAGTGAAGCCACAGATGTATAAACATTAATGAACACACTTATGCCTACTGCCTCGGA] and sh-2 [5’ TGCTGT TGACAGTGAGCGACCGGGACAACGTTTATTACTATAGTGAAGCCACAGATGTATAGTAATAAACGTTGTCCCGGGTGCCTACTGCCTCGGA]) short-hairpin RNAs (shRNA). Cells were selected with growth media supplemented with 1ug/ml Puromycin (Life #A11138-03).

### Plasmid DNA overexpression

MDA-MB-231 cells were stably transfected using FUGENE6 with empty or hE-cadherin-pcDNA3 vector (Addgene #45769) using 15ug DNA per 10cm plate of cells. Cells were selected in growth media supplemented with 800ug/ml G418 (Invitrogen #10131-035).

### Immunoblotting

Samples for immunoblot analysis were collected using RIPA buffer (50mM Tris pH 7.4, 150mM NaCI, 1mM EDTA, 0.5% Nonidet P-40, 0.5% NaDeoxycholate, 0.1% SDS, 1x HALT cocktail [Thermo Fisher #78442]) and standard immunoblot technique was followed. Membranes were blocked in Odyssey PBS Blocking Buffer (LiCor #927-40000), and incubated in primary antibodies overnight: plGF1RY1135 (Cell Signaling #3918; 1:500), IGF1R β-subunit (Cell I Signaling #3027; 1:1000), pAkt S473 (Cell Signaling #4060; 1:1000), total Akt (Cell Signaling i #9272; 1:1000), E-cadherin (BD Biosciences #610182; 1:1000), and β-actin (Sigma #A5441; 1:5000). Membranes were incubated in LiCor secondary antibodies for 1 hour (anti-rabbit 800CW [LiCor #926-32211]; anti-mouse 680LT [LiCor #925-68020]; 1:10,000), and imaged with Odyssey Infrared Imager.

### IGF1-induced cell cycle and viability analysis

*For cell cycle:* MCF-7 and ZR75.1 cells were reverse transfected as described above, serum starved for approx. 30 hours, and pulsed with 10nM IGF1 for 17 hours. Cells were fixed in 70% EtOH for 30 minutes at 4°C and RNA digested using 50ng/ul RNase A (Qiagen #1007885) for 15 minutes at 37°C. DNA content was then stained using 50ng/ul propidium iodide (Sigma #P4170) for 30 minutes at 4°C. Cell cycle profiles were analyzed using the BD LSRII flow cytometer and analyzed using the FACS DIVA software. The statistical difference in percent of cells in S- or G2/M phase in IGF1 treated cells over vehicle control in experimental groups was evaluated using a two-tailed student’s t-test (p<0.05).

*For viability:* T47D shSCR and shCDH1 #1 and #2 cells were plated in serum-free media in 96 well plates (9,000 cells/well) and then stimulated with IGF1 (10nM) for 6 days. The FluoReporter Blue Fluorometric dsDNA Quantitation Kit was used to measure DNA content. Statistical difference in Hoechst fluorescence in IGF1 treated cells over vehicle control in each cell line was evaluated using a two-tailed student’s t-test (p<0.05).

### Immunofluorescence and Proximity lipation assay (PLA)

Cells were plated on coverslips and fixed in 4% paraformaldehyde for 30 minutes at 37°C. Coverslips were permeabilized for 1 hour using PBS+0.3% Triton X-100. for immunofluorescence, coverslips were blocked in PBS+5% goat serum, incubated in primary antibody overnight (total IGF1R β-subunit [Cell Signaling #3027; 1:300] and E-cadherin [BD Biosciences #610182; 1:100]), followed by Alexa Fluor secondary antibody incubation for 1 hour (anti-rabbit Alexa Fluor 488 [Life Technologies #A11070] and anti-mouse Alexa Fluor 546 [Life Technologies #A11018]; 1:200). For *in situ* proximity ligation assay, coverslips were processed using the Duolink Red mouse/rabbit kit using the protocol provided (Sigma #DUO92101) with the antibody dilutions above. The ratio of puncta/nuclei for each experimental condition was calculated by counting all puncta and nuclei in five 60x images. One-way ANOVA was used to compare the ratios between the experimental conditions (VHC, 30m, 6hr, 24hr). Confocal microscopy was used for imaging.

### Dose response growth assays and synergy measurements

MCF-7 and ZR75.1 cells were reverse transfected with control or CDH1 siRNA as described above into 96-well plates (9,000 cells/well) in 100ul of media/well. Cells were treated with 3x vehicle (DMSO), OSI-906 (Selleckchem #S1091) or BMS-754807 diluted in 50ul of media for a final volume in each well of 150ul (n=6 per concentration). Plates (2D and ultra-low attachment [ULA; Corning #3474]) were collected on day 6 and viability was measured using CellTiter Glo Viability assay (Promega #G7572). EC_50_ values for viability were calculated by non-linear regression and statistical differences evaluated using sum-of-squares Global f-test (p<0.05). For synergy experiments, SUM44PE and MDA-MB-134 cells were plated in 96-well ULA plates (18,000 cells/well) in 100ul of media/well. Cells were treated with 6x vehicle (DMSO), OSI-906, BMS-754807, or BEZ235 (Selleckchem #S1009) diluted in 25ul of media such that the combination of two drugs resulted in 150ul of total volume in each well (n=2 per experiment). Synergy was calculated using the Median-Effect Principle and Combination Index-Isobologram Theorem (Chou-Talalay)^27^. Combination index values for ED50, ED75, ED90 are shown as a mean ± SEM from n=3 independent experiments.

### In vivo ILC xenograft growth and explant culturing

MDA-MB-134 cells (5x10^6^ cells) and BCK4 cells (5x10^6^ cells) were injected into the right inguinal mammary fat pads of 7-8 week old NOD.Cg-Prkdcscid Il2rgtm1Wjl/SzJ (NSG; The Jackson Laboratory) and NOD.CB17-Prdkcscid/J mice (NOD SCID; The Jackson Laboratory), respectively (implanted with 0.36mg 90-day slow release estradiol pellets [Innovative Research of America #SE-121]) and grown to a tumor volume of 350mm^3^. Tumors were collected, minced into 1-2mm^3^ chunks of tumor tissue, and plated onto Vetspon Absorbable Hemostatic Gelatin sponges (Patterson Veterinary #07-849-4032) in 12-well tissue culture plates containing 1.5mls of explant media (DMEM/F12+10% FBS with 10mM HEPES, 1mg/ml BSA, 10ug/ml insulin, 10ug/ml hydrocortisone, 1x antibiotic-antimycotic solution [Thermo Fisher #15240-062]). Media was treated with vehicle or 1uM BMS-754807 for 72 hours. Tissue was collected by formalin fixation followed by paraffin embedding. Sections were stained for Ki67 (Dako #M7240; 1:100) using standard immunohistochemistry technique. Nuclei were quantified by counting all clearly defined nuclei within each tissue section (n=3-6). Two-tailed student’s t-test was used to determine statistical difference between vehicle and BMS-754807 treatment (p<0.05).

### TCGA Data Analysis

TCGA RNA-seq expression data were downloaded as transcripts per million (TPM) from the Gene Expression Omnibus database (GEO: GSE62944) and log2(TPM+1) for gene-level results were used. TCGA Reverse Phase Protein Array (RPPA) data were downloaded as median-normalized, batch-corrected expression values from TCPA (Level 4, version 4.0). ER+ IDC (n=417) and ILC (n=137) samples with both RNA-Seq and RPPA data were used for all analyses. Mann-Whitney U tests were used to compare expression, Spearman’s rho to compare correlations, and a chi-square test to compare proportions between ILC and IDC tumors. All were calculated using R (version 3.4.1). The median expression values for IGF1 and pIGF1R across ER+ IDC and ILC tumors (n=554) were used as cutoffs for Figure 4G.

## RESULTS

### Loss or inhibition of E-cadherin results in enhanced IGF1R activity

To validate our previously published data^17^ and to further understand the regulation of the IGF1 signaling pathway by E-cadherin, we silenced E-cadherin *(CDH1)* by siRNA knockdown in a panel of three estrogen receptor (ER)-positive IDC cell lines and then stimulated with a dose series of IGF1 (0, 1,10, 100nM). MCF-7, ZR75.1, and T47D E-cadherin knockdown (siCDH1) cells showed enhanced sensitivity to IGF1 compared to the scramble control (siSCR) cells, most notable at the 1nM dose of IGF1, resulting in increased phosphorylation of IGF1R and Akt (Fig 1A-C). As a complementary approach, we inhibited E-cadherin function in MCF-7 cells using the HECD-1 monoclonal antibody that binds the extracellular domain of E-cadherin and prevents adherens junction formation. Similar to the knockdown of E-cadherin, HECD-1 treated cells showed increased IGF1R and Akt phosphorylation compared to control (Fig 1D). Additionally, we evaluated confluency-dependent IGF1R signaling to understand the effect of increased cell-cell contacts. A confluent monolayer of MCF-7 cells lost the ability to initiate IGF1R signaling upon ligand stimulation compared to a sub-confluent monolayer (approx. 40-50%), however, the knockdown of E-cadherin rescued signaling in both confluency conditions (Fig 1E).

**Figure 1:**
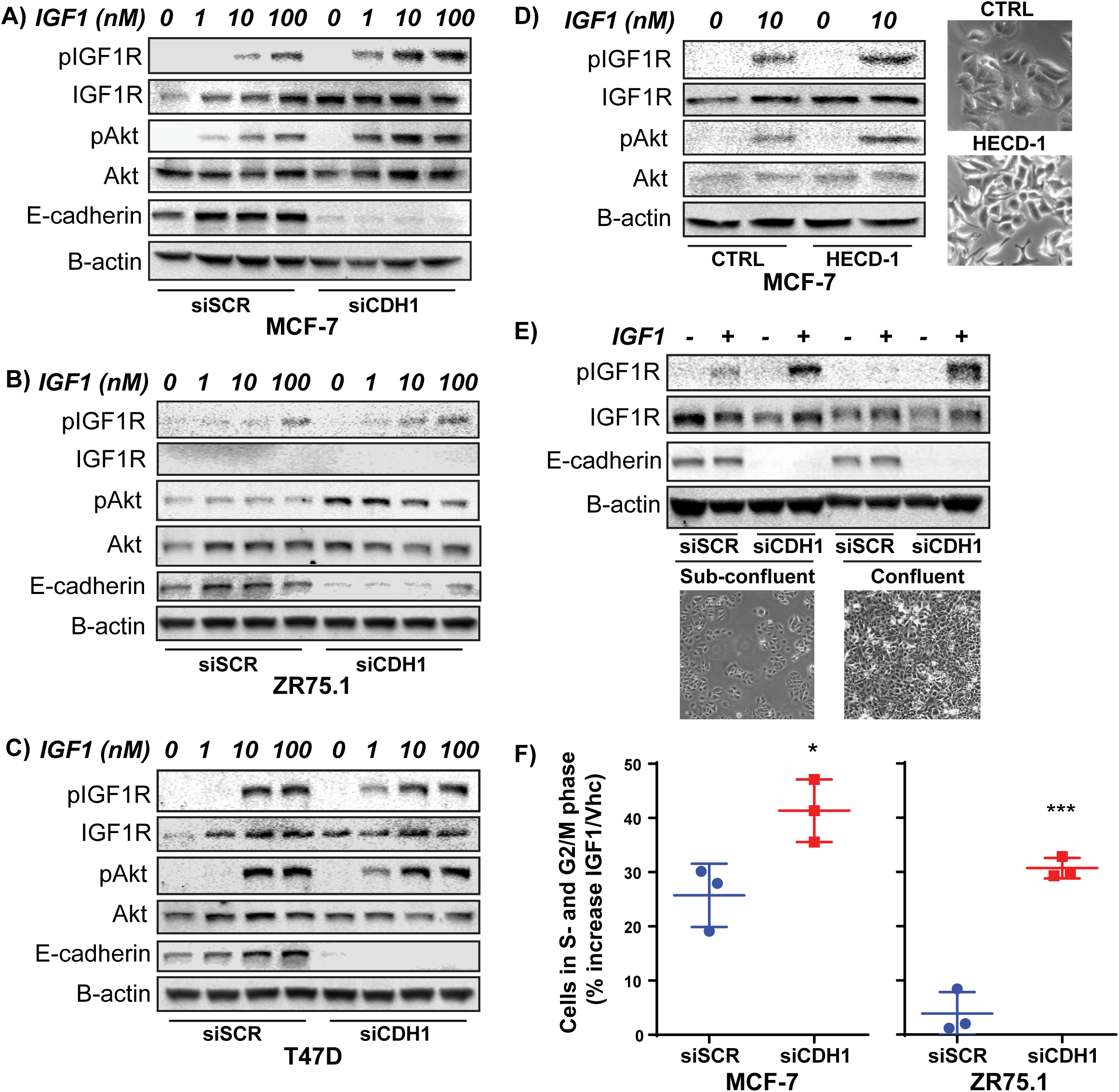
Loss or inhibition of E-cadherin *(CDH1)* expression enhances IGF1R signaling. (A) MCF-7, (B) ZR75.1, and (C) T47D breast cancer cells transfected with SCR (siSCR) or CDH1 (siCDH1) siRNA were stimulated with increasing doses of IGF1 (0-100nM) for 10 min. IGF1R and Akt signaling was assessed by immunoblot. Of note, IGF1R expression could routinely not be detected in ZR75.1. (D) MCF-7 cells were treated with 25ug/ml HECD-1 antibody for 24 hours and imaged by phase-contrast microscopy for dissociation of adherens junctions. Cells were stimulated with Vhc or 10nM IGF1 for 10 min and IGF1R and Akt signaling assessed by immunoblot. (E) MCF-7 cells were plated at sub-confluency (200k cells in 6-well) or high confluency (800k cells) and then stimulated with either Vhc or 10nM IGF1 for 10 min. IGF1R signaling was assessed by immunoblot. Representative phase-contrast microscopy images of the cell plating densities are shown. (F) MCF-7 and ZR75.1 siSCR and siCDHI cells were serum-starved and stimulated with 10nM IGF1 for 17 hours and DNA stained with propidium iodide to measure cell cycle profile. The percent of cells in the IGF1 A/hc conditions in the S- and G2/M phases of the cell cycle for siSCR and siCDHI are shown (representative experiment shown; n=2 or 3 each with 3 biological replicates).

We evaluated the functional effect of enhanced IGF1 signaling on the cell cycle profile in MCF-7 and ZR75.1 cells with reduced E-cadherin. CDH1 knockdown cells showed a significant increase (p=0.03 and p=0.0005, respectively) in the percentage of cells progressing into the S- and G2/M-phases of the cell cycle following IGF1 treatment compared to siSCR cells (Fig 1F). Similarly, slight increases in IGF1-induced cell viability in siCDH1 compared to siSCR in T47D cells were observed (Fig S1).

We overexpressed E-cadherin in MDA-MB-231 cells, an ER-negative IDC cell line with undetectable E-cadherin protein by immunoblot to determine if overexpression represses signaling. Although adherens junction formation was not observed (data not shown), E-cadherin overexpressing cells demonstrated decreased phosphorylation of IGF1R and Akt compared to empty vector control cells, and significantly less cell cycle progression in response to IGF1 stimulation (p=0.011; Fig S2).

### Loss of E-cadherin enhances sensitivity to IGF1R inhibition

Due to the enhanced sensitivity of E-cadherin knockdown cells to IGF1 stimulation, we determined if loss of E-cadherin in MCF-7 and ZR75.1 cells also increased sensitivity to the IGF1R ATP-competitive small molecule inhibitors, OSI-906 (OSI) and BMS-754807 (BMS). In addition to 2D adherent culture, ultra-low attachment suspension growth (ULA) was examined, since we observed increased cell viability in E-cadherin knockdown cells under these conditions (Tasdemir et al, manuscript in preparation), possibly due to the reported annoikis resistance of cells lacking E-cadherin expression^28^. MCF-7 siCDH1 cells displayed significantly decreased viability in response to OSI treatment, compared to siSCR cells in both 2D (p<0.0001; Fig 2A) and ULA (p=0.0003; Fig 2B) growth conditions resulting in a shift in the EC_50_. Additionally, ZR75.1 siCDH1 cells showed significantly decreased viability and a shift in the EC_50_ when grown in ULA (p<0.0001; Fig S3) in response to OSI treatment, but not in the 2D growth condition. Similarly, MCF-7 siCDH1 cells showed decreased viability in response to BMS compared to siSCR cells the ULA growth condition (p<0.0001), but no significant difference in 2D (Fig 2C-D). Overall, these data suggest that the loss of E-cadherin enhances breast cancer cell sensitivity to IGF1R inhibition. We also tested the growth response of MCF-7 siSCR and siCDH1 cells treated with ICI 182,780 (ICI), a selective estrogen receptor downregulator (SERD), and observed no statistical difference in EC_50_ suggesting that the loss of E-cadherin does not generally sensitize cells to all small molecule drug treatments (Fig S4).

**Figure 2:**
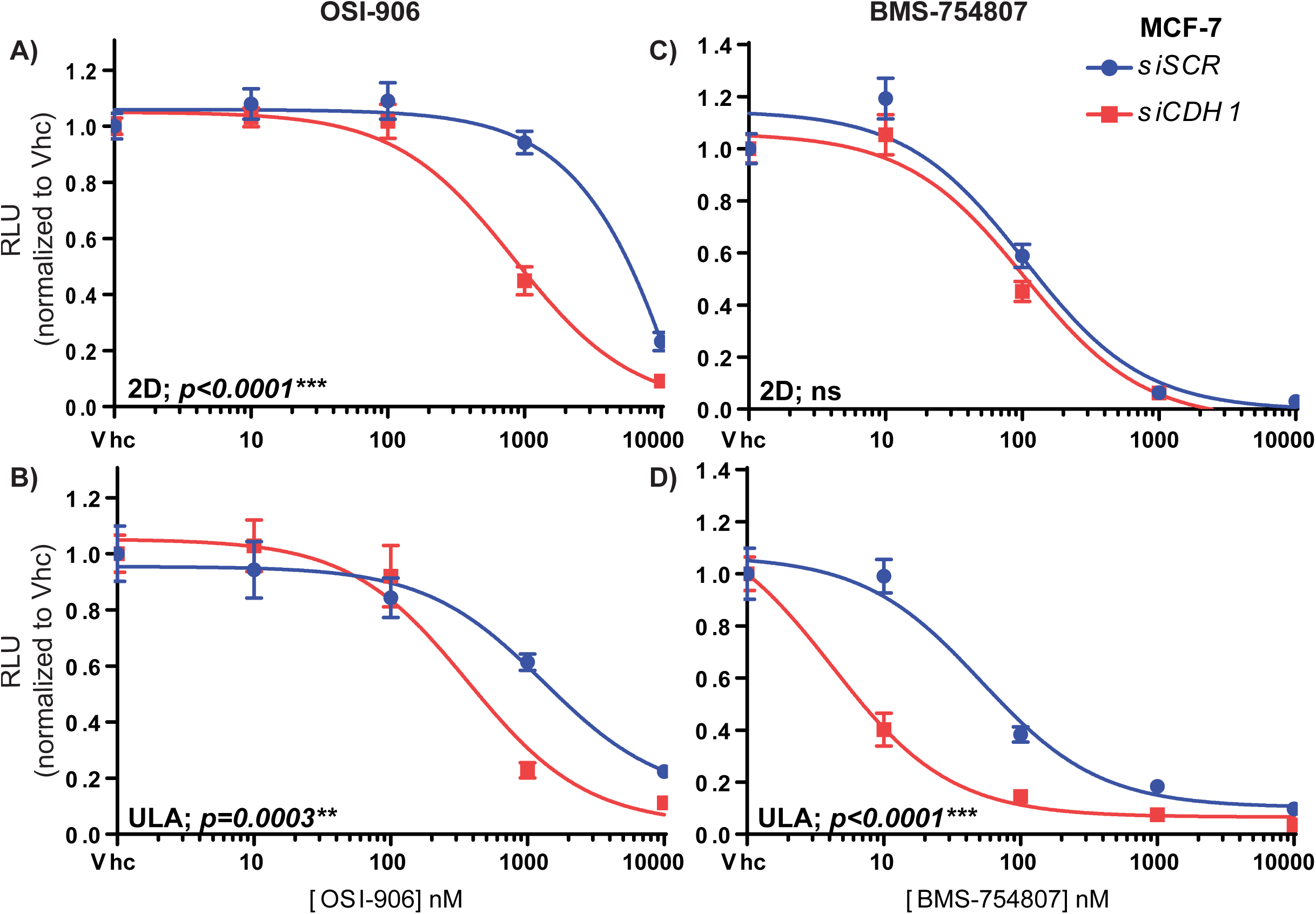
Knockdown of E-cadherin increases sensitivity to IGF1R inhibition in breast cancer cells. MCF-7 cells were reverse transfected with SCR or CDH1 siRNA and seeded into 96-well 2D or ULA plates and treated with IGF1R inhibitor (OSI-906 or BMS-754807) for 6 days. Conditions in the panels as follows: (A) OSI-906; 2D, (B) OSI-906; ULA, (C) BMS-754807; 2D, (D) BMS-754807; ULA. The CellTiter Glo assay was used to assess cell viability (relative luminescence). EC50 values for viability were calculated by non-linear regression and statistical differences evaluated using sum-of-squares Global f-test (p<0.05; representative experiment shown; n=3 each with 6 biological replicates).

### IGF1R and E-cadherin directly interact in ER+ breast cancer cells resulting in recruitment of IGF1R to adherens junctions

To understand how E-cadherin regulates IGF1R, we assessed whether IGF1R and E-cadherin directly interact in breast cancer cells using *in situ* proximity ligation assay (PLA). The sensitivity and specificity of PLA allows for detection of endogenous interacting proteins within proximity of no further than 40nm. PLA showed that IGF1R and E-cadherin directly interact in both MCF-7 and T47D cells, as shown by the red fluorescent puncta (Fig 3A-B). To demonstrate the specificity of the detection, we used MCF-7 knockdown cells lacking E-cadherin (siCDH1) or IGF1R (siIGFR) as negative controls and observed the red fluorescent puncta signal greatly diminished (Fig 3C-D, Fig S5A-C). Additionally, secondary antibody specificity was confirmed by using each primary antibody alone and a no primary antibody control and did not detect significant levels of PLA puncta over background (Fig S5D-F). The interaction between IGF1R and E-cadherin following IGF1 stimulation was examined using PLA. In MCF-7 cells, IGF1 treatment caused a significant decrease in number of fluorescent puncta (p=0.003), suggesting that the interaction between the two proteins needs to be disrupted for proper IGF1R function (Fig 3E-I), possibly explaining why siCDH1 cells have an increased IGF1R signaling capacity compared to control cells.

**Figure 3:**
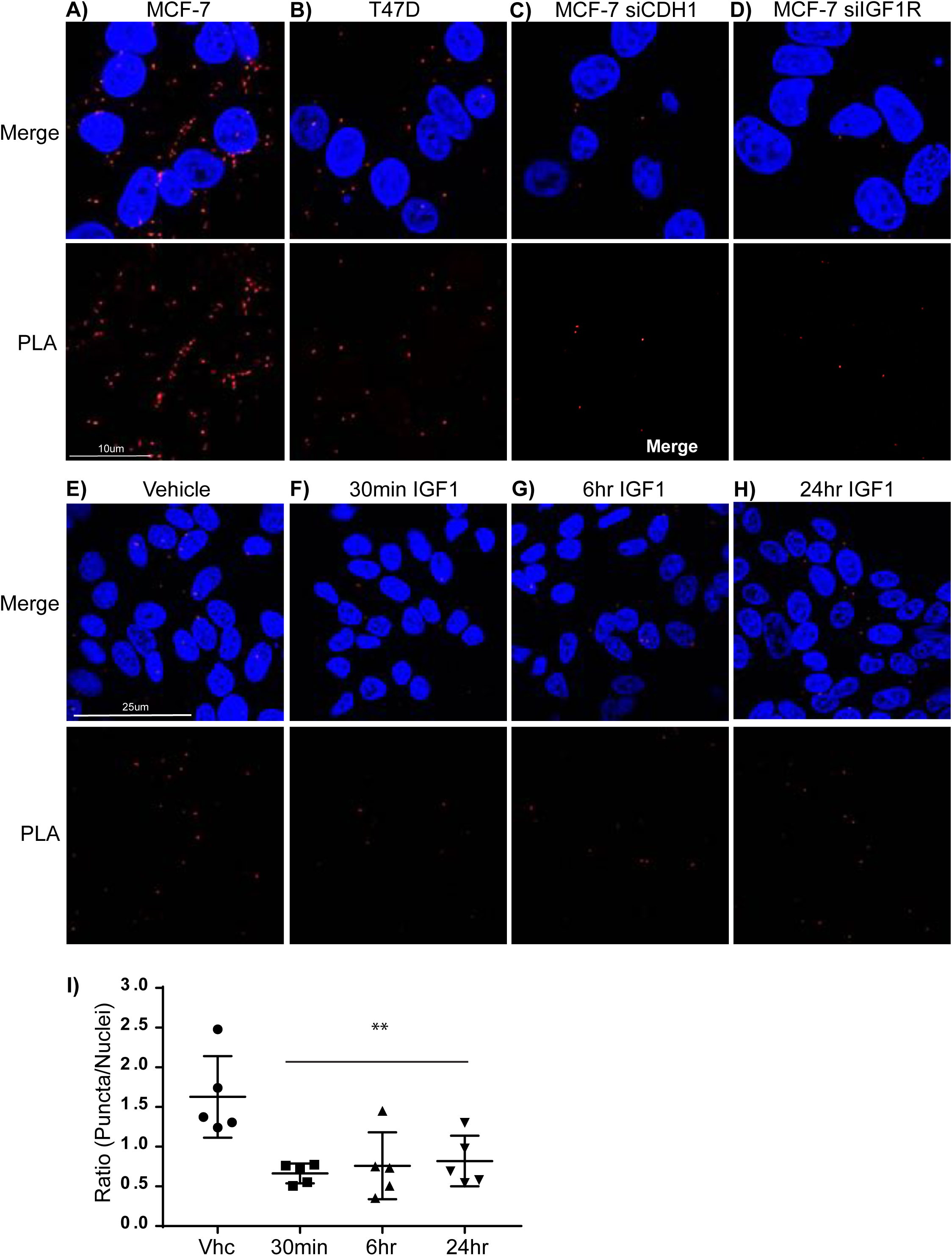
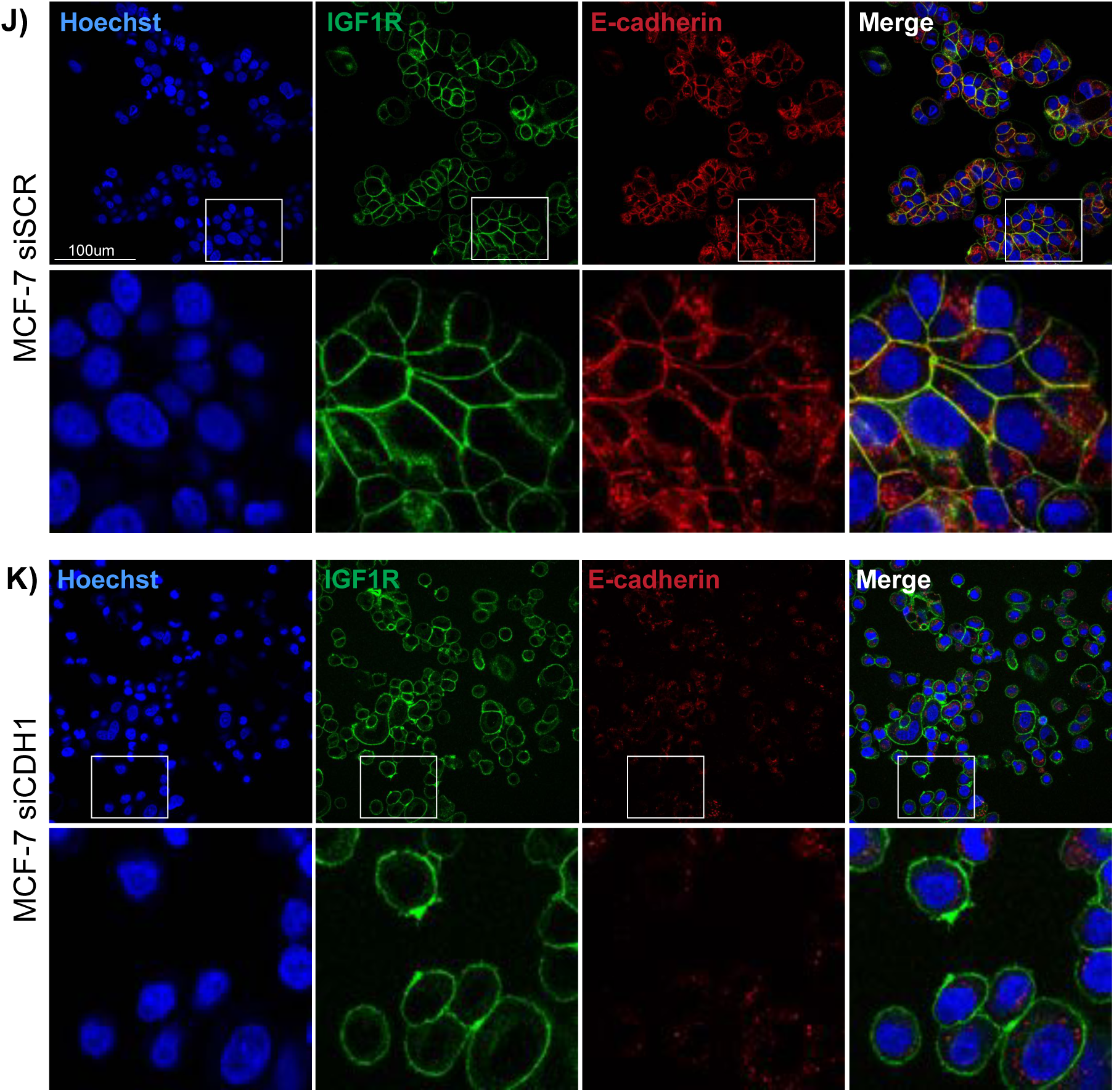
Proximity ligation assay reveals direct interaction between IGF1R and E-cadherin and recruitment of IG1R to adherens junctions. *In situ* proximity ligation assay (PLA) was used to analyze the direct interaction between IGF1R and E-cadherin in breast cancer cells. (A) MCF-7 and (B) T47D cells were plated on coverslips, fixed, and stained with IGF1R and E-cadherin antibody overnight. The Duolink (Sigma) protocol was followed and coverslips were imaged using confocal microscopy to reveal red puncta. (C) MCF-7 siCDHI and (D) silGF1R cells were used as negative controls for the assay to assess primary antibody specificity. MCF-7 cells were plated on coverslips and treated with either (E) Vhc or 10nM IGF1 for (F) 30 minutes, (G) 6 hours, or (H) 24 hours. PLA protocol for IGF1R and E-cadherin was followed as described above. (I) Red puncta and nuclei (stained with DAPI) were quantified and displayed as a ratio of puncta/nuclei. All puncta and nuclei in 60x images were counted. Oneway ANOVA was used to determine significant difference between groups (p<0.05; one independent experiment, n=5 images per slide counted). The co-localization of IGF1R (green) and E-cadherin (red) was analyzed by immunofluorescence staining in (J) MCF-7 siSCR and (K) siCDHI knockdown cells.

**Figure 4:**
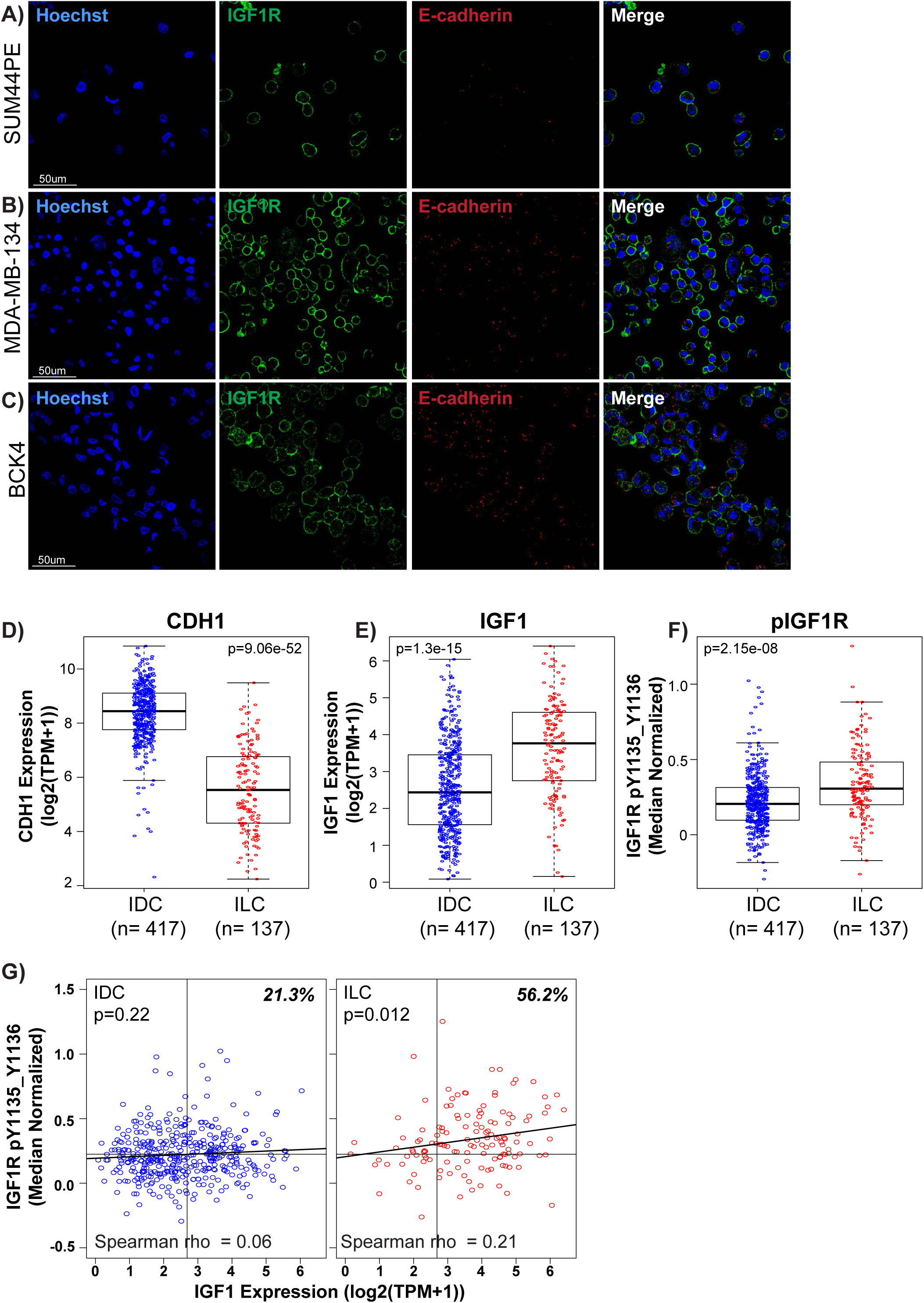
IGF1-IGF1R pathway is active in invasive lobular breast carcinoma with genetic loss of *CDH1*. (A) SUM44PE, (B) MDA-MB-134, and (C) BCK4 ILC cells were immunostained for IGF1R (green) and E-cadherin (red) and imaged by confocal microscopy. Of note, BCK4 cells were imaged at an increased exposure compared to MM134 and SUM44PE cells. (D) CDH1 mRNA, (E) IGF1 mRNA, (F) and plGF1 R Y1135 & Y1136 levels in ER+ IDC compared to ER+ ILC in TCGA were plotted using RNAseq (Iog2 TPM+1) and RPPA (median normalized) data. The TCGA cohort includes n=417 IDC cases and n=137 ILC cases that have matched data for RNAseq and RPPA. Man-Whitney test was used to determine significant differences in expression level between the two subtypes, p<0.05). (G) Correlation between pIGFIR and IGF1 ligand expression is plotted for IDC (left) and ILC (right). Spearman’s rank correlation was used to demonstrate the correlation between the two variables with significance as defined by p<0.05.

We stained MCF-7 cells for endogenous IGF1R and E-cadherin and determined that IGF1R and E-cadherin co-localize to adherens junctions. Interestingly, co-localization was prominent at the points of cell-cell contact, and noticeably absent or reduced on portions of the membrane where there was no cell-cell contact (Fig 3J). This suggests that E-cadherin recruits IGF1R to adherens junctions, perhaps to sequester the receptor as a mechanism of signaling repression. Upon knockdown of E-cadherin the expression pattern of IGF1R appears to redistribute equally to the entire cell membrane (Fig 3K) supporting the idea that E-cadherin influences and regulates IGF1R localization.

### Invasive lobular breast cancers (ILC) display enhanced IGF1-IGF1R pathway activation

Because knockdown or inhibition of E-cadherin induces hyperactivity of the IGF1R pathway in cell line models, we investigated whether IGF1R pathway activity is also hyperactivated in ILC, a subtype of breast cancer that accounts for 10-15% of all breast cancer cases and is molecularly classified by its genetic loss of E-cadherin^23^. Because 90-95% of ILC tumors are ER+, we focused on this cohort^23^. IGF1R expression and localization was examined in the ER+ ILC cell lines: MDA-MB-134 (MM134; Fig 4A), SUM44PE (Fig 4B), and BCK4 (Fig 4C). IGF1R staining was membranous similar to that observed in MCF-7 siCDH1 cells (Fig 3F). As expected, ILC cells showed a lack of membranous E-cadherin staining (Fig 4A-C).

To compare IGF1R activity in ER+ ILC and IDC tumors, CDH1 and IGF1 ligand mRNA expression, and IGF1R phosphorylation (pIGF1R; Y1135/Y1136) were examined using RNA-sequencing and Reverse Phase Protein Array data from The Cancer Genome Atlas (TCGA). Concurrent with a decrease in CDH1 mRNA expression (p=9.06e-52; Fig 4D), IGF1 ligand mRNA expression (p=1.3e-15; Fig 4E) and pIGF1R levels (p=2.15e-08; Fig 4F) were significantly increased in the ILC tumors compared to IDC tumors. Interestingly, ILC tumors exhibited a significant positive correlation between IGF1 mRNA expression and pIGF1R level (Spearman rho=0.21; p=0.012), despite having significantly reduced total IGF1R expression compared to IDC (data not shown; Fig 4G). In contrast, IDC tumors did not show a correlation (Spearman rho=0.06; p=0.22) suggesting that presence of IGF1 ligand did not necessarily activate IGF1R in IDC. Strikingly, the percentage of tumors with higher than median expression (across all breast tumors) of both IGF1 and pIGF1R is significantly higher in ILC (56.2%) compared to IDC (21.3%), suggesting that IGF1 ligand activates IGF1R signaling in these tumors more efficiently with the loss of E-cadherin (chi-square test, p= 2.5e-14 [Fig 4G]). Interestingly, when assessing activation of the IGF-sig^16^ in ER+ ILC versus IDC in the TCGA cohort we did not observe a difference in expression score (data not shown).

### IGF1R inhibitors and endocrine therapy synergize to decrease viability in ILC cells

Clinically, patients with ER+ ILC are treated with endocrine therapy targeting ER, however, data from the BIG 1-98 trial suggest that ILC tumors demonstrate resistance to tamoxifen, a selective estrogen receptor modulator, compared to IDC^29^. Additionally, results from multiple clinical studies indicate that ILC patients have a poorer prognosis with more frequent late recurrences compared to IDC^30–32^. This highlights the need to improve therapeutic options in ILC patients based on uniquely activated pathways and therefore, we evaluated efficacy of IGF1R pathway inhibitors in ER+ ILC cell lines in combination with endocrine therapy. Recent data published from our lab suggest that tamoxifen, can act as a partial ER agonist activating ER activity in some ILC cell lines, rather than a pure antagonist as in IDC cells^33^, in line with the data from the BIG1-98 study. Therefore, we tested efficacy of the selective estrogen receptor downregulator, ICI 182,780 (ICI) in combination with two IGF1R inhibitors used in Figure 2 (OSI and BMS) and a PI3K/mTOR inhibitor (BEZ235 [BEZ]). SUM44PE and MM134 cells were treated with increasing doses of OSI (Fig 5A-B; Fig S6A-B), BMS (Fig 5C-D; Fig S6C-D), and BEZ (Fig 5E-F; Fig S6E-F) in combination with increasing doses of ICI. With all three IGF1R pathway inhibitors, decreased cell viability was observed with the addition of increasing doses of ICI. Formal synergy testing of the drug combinations using the Median-Effect Principle and Combination-Index Isobologram Theorem, commonly referred to as the Chou-Talalay method^27^ revealed combination index (CI) values less than 1 for drug interactions at the ED50, ED75, and ED90 indicating a high level of synergy for the three sets of inhibitor combinations (Fig 5, Fig S6, Table S1). The lowest CI values were observed for the BMS+ICI drug combination in SUM44PE cells (ED50=0.127, ED75=0.081, ED90=0.099). Additionally, a minimum dose reduction index (DRI) for ICI of 8-fold for all drug combinations in SUM44PE cells and 2-fold in MM134 cells at the EC50 was seen. This data suggests that adding an IGF1R pathway inhibitor in combination with ICI reduces the concentration of ICI necessary to achieve that same inhibitory effect as ICI alone.

**Figure 5:**
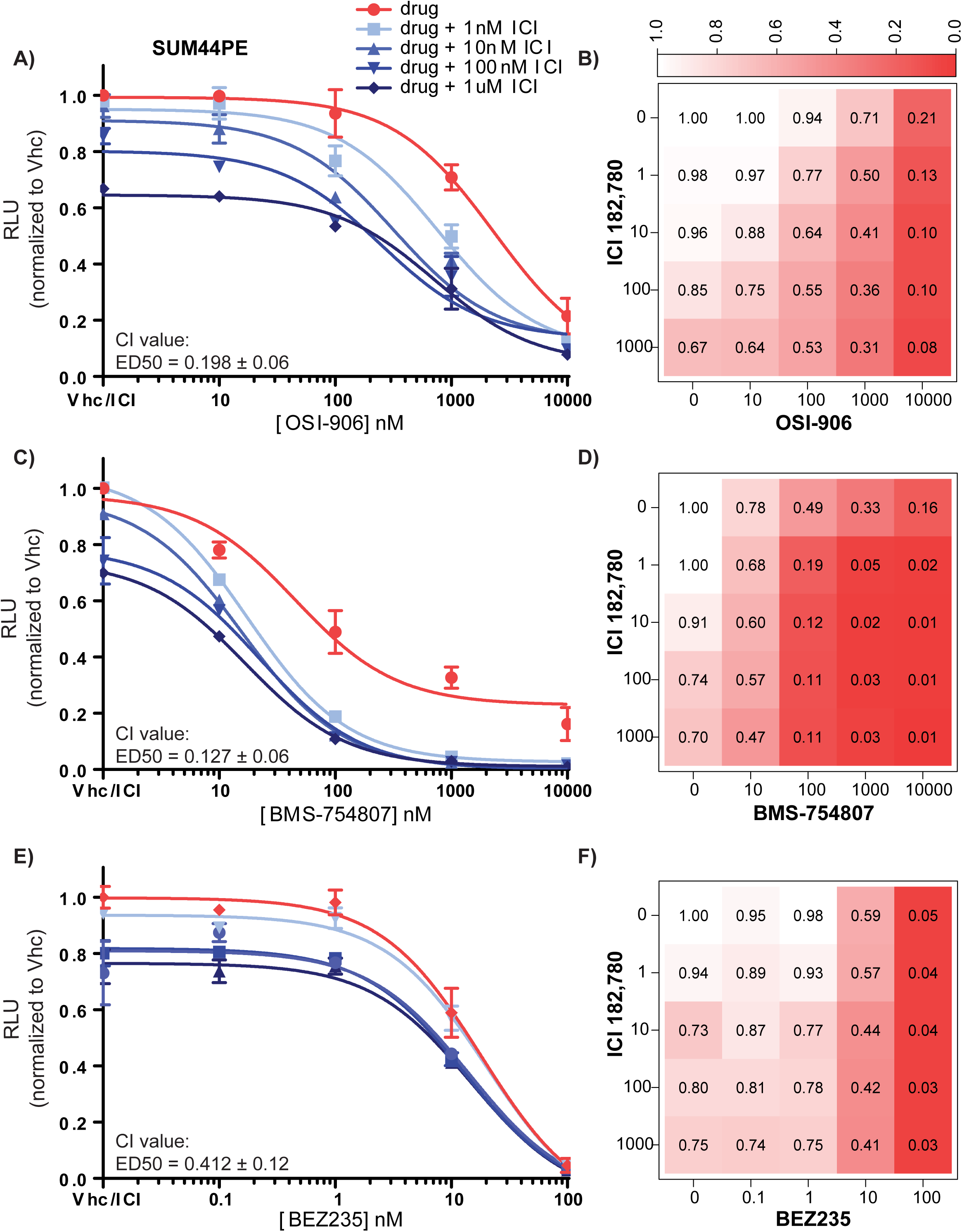
IGF1R pathway inhibitors and endocrine therapy synergize to inhibit cell viability in ILC breast cancer cells. SUM44PE ILC cells were plated into 96-well ULA plates and treated for 6 days with increasing doses of (A, B) OSI-906, (C, D) BMS-754807, or (E, F) BEZ235 in combination with increasing doses of ICI 182,780. The dose response curves and heat maps shown indicate inhibition of cell viability (CellTiter Glo). Representative experiment shown; n=3 independent experiments each with 2 biological replicates per combination of doses.

### Ex vivo IGF1R inhibition inhibits proliferation in an ILC xenograft

Finally, we evaluated the efficacy of an IGF1R inhibitor in ILC tumors. However, there are a limited number of ILC patient-derived xenograft (PDX) and cell line xenograft models, and their slow growth rates makes large scale *in vivo* studies challenging. We therefore treated two ILC cell line xenografts *ex vivo* as explant cultures, as previously described^34–37^. The advantages of this technique include less tissue requirement for the assay compared to an *in vivo* study and rapid understanding of the therapeutic efficacy of the inhibitor. Additionally, data published by Majumder et al.^37^ suggest a high concordance between *ex vivo* and *in vivo* tumor response to drug treatment. MM134 and BCK4 cells (a weakly ER responsive ILC cell line, not used for synergy experiments due to slow growth *in vitro)* were grown as xenografts, harvested and plated as explant culture, and treated with vehicle or BMS (1μM) for 72 hours. The tissue was collected and stained for Ki67 as a marker of proliferation. We observed a significant decrease in Ki67 positive nuclei in both tumor models treated with BMS (Fig 6). In the MM134 tumor we observed a significant decrease (p=0.002) in Ki67 positive nuclei from 47% in the vehicle to 22% in the BMS treated tumor tissue (n=3 or 4; Fig 6A-C). Similarly, in the BCK4 tumor we observed a significant decrease (p=0.005) in Ki67 positive nuclei from 25% in the vehicle to 11% in the BMS treated tumor tissue (n=6; Fig 6D-F). This data suggest that targeting IGF1R in ILC tumors may be a useful strategy to inhibit cell proliferation.

**Figure 6:**
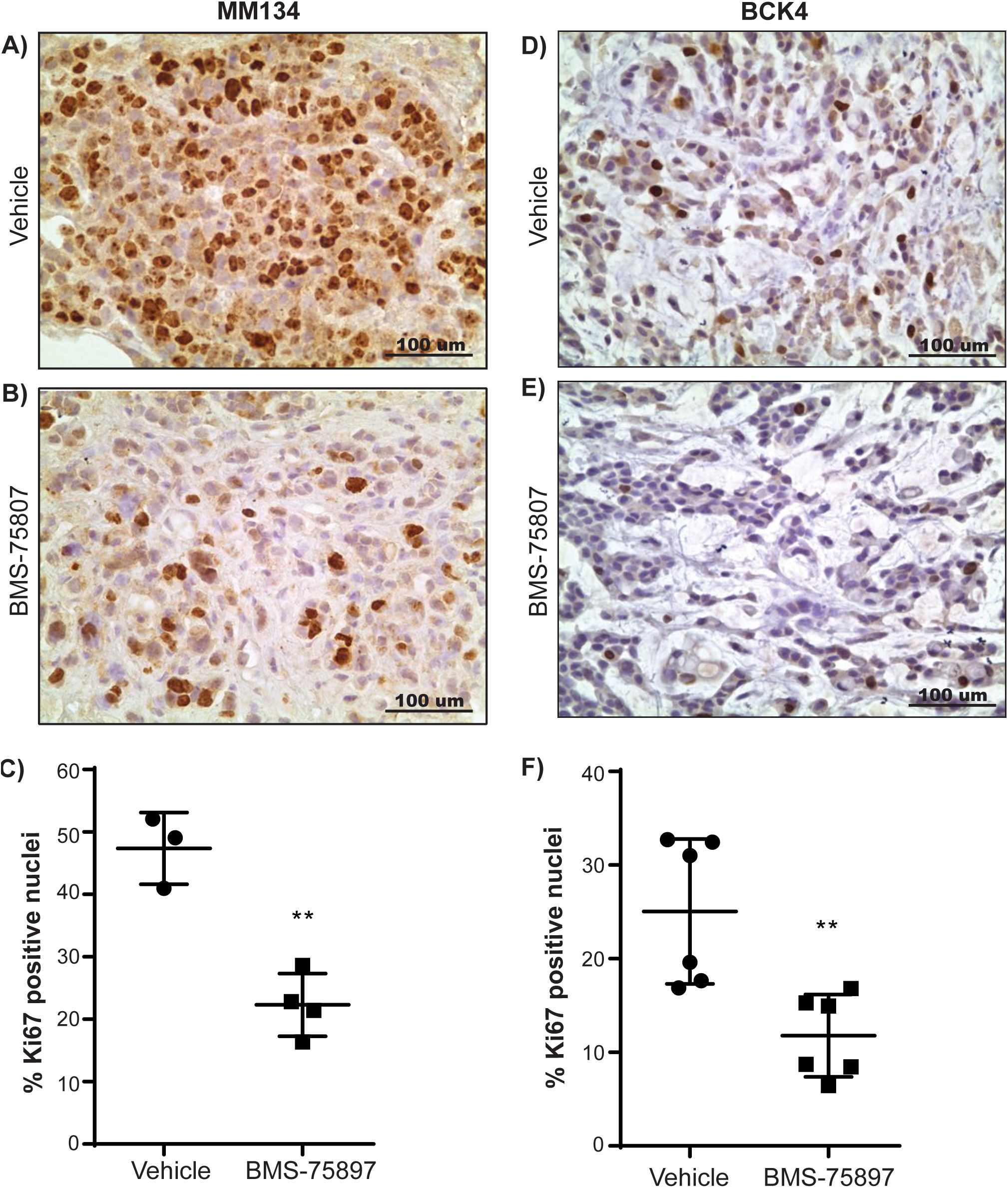
IGF1R inhibition reduces Ki67 staining in ILC tumor *ex vivo* culture. MM134 and BCK4 xenograft tumors were harvested from immunocompromised mice, minced into 1-2mm^3^ tumor chunks and then plated on gelatin sponges in 12-well plate containing 1.5ml media. Media was treated with DMSO Vhc or 1uM BMS-75807 for 72 hours. Tumor pieces were harvested by FFPE and stained for Ki67 as a marker of proliferation (A-B, MM134; D-E, BCK4). Staining was quantified by counting all clearly defined nuclei in 20x images (C and F). Statistical difference was assessed using a Student’s t-test (p<0.05; n=3-6).

## DISCUSSION

Despite a large body of preclinical evidence supporting the use of IGF1R inhibitors for the treatment of breast cancer, the outcomes of clinical trials testing the efficacy of these drugs in patients thus far have been disappointing. However, these trials proceeded with a lack of appropriate biomarkers for predicting positive therapeutic efficacy and little to no understanding of which tumor types would benefit^13–15,38^. In response, in recent years the field has emphasized the need to understand and identify gene expression or proteomic biomarkers that predict a positive response to targeted therapy. Along this thought process, we previously published a gene expression signature used to identify tumors that are IGF1 responsive^16^ and here we focus on one proteomic biomarker, E-cadherin, identified through an integrative computational approach recently published by our group^17^. It is known that constitutive IGF1R activation drives E-cadherin transcriptional repression through EMT^6,10^, however, the reverse regulation of IGF1R by E-cadherin has not been previously characterized. Our data suggest that loss of E-cadherin in breast tumors, specifically in ILC, highlights a subset of tumors that may be responsive to IGF1R inhibition and here we begin to describe the mechanism by which this regulation occurs.

We demonstrate that in breast cancer cells, IGF1R is endogenously localized to cell-cell contacts, similar to data published in MCF-7 cells overexpressing IGF1R^39^ and in corneal epithelial cells^40^. We show a direct, endogenous interaction between IGF1R and E-cadherin using *in situ* proximity ligation assay. To our knowledge interaction between IGF1R and E-cadherin in breast cancer cells has only been demonstrated by immunoprecipitation (IP)^39^. Our data provide confirmation of this interaction using a technique known to be higher in specificity and sensitivity compared to IP, which requires intense cell manipulation (e.g. lysis and scraping) and often results in pull-down of entire protein complexes. This suggests that IGF1R is recruited to adherens junctions by E-cadherin, possibly resulting in receptor sequestration and signaling repression. This process is similar to the sequestration of EGFR into the adherens junction and loss of receptor mobility, a well characterized mechanism of EGFR signaling repression^41–43^. However, data published by Curto et al. suggests that the latter action is mediated through the tumor suppressor, Merlin, responsible for coordinating stabilization of the adherens junction and thereby regulating contact-inhibition growth^43^. Although IGF1R signaling is controlled in a contact-dependent manner (Fig 1E), they also showed that IGF1 activity is not regulated by Merlin, indicating that IGF1R regulation by E-cadherin likely occurs independent of this factor^43^. Although there may be a yet undefined intermediate regulator similar to Merlin, our data indicate that E-cadherin plays a role in coordinating the recruitment and sequestration of IGF1R within the adherens junction to repress IGF1R signaling. When E-cadherin is lost and junction formation is disrupted (such as in ILC cells), IGF1R is released and re-localizes to the entirety of the cell membrane where signaling is more easily initiated upon IGF1 ligand binding.

Supporting this concept, our data indicate that the knockdown of E-cadherin in three ER+ breast cancer cell lines not only enhanced IGF1-induced signaling via IGF1R but also increased sensitivity of the cells to the ligand. This is similar to the relationship reported between EGF-EGFR and IGF1-IGF1R upon adherens junction disruption via calcium-depletion^41^. Because of the increased IGF1R pathway activation associated with the loss of E-cadherin, the knockdown cells in turn became more sensitive to IGF1R inhibition.

We believe that IGF1R signaling may be particularly important in ILC, an understudied subtype of breast cancer, due to the complete loss of E-cadherin protein and/or adherens junction formation. In this subtype, the loss of E-cadherin may serve as a biomarker of IGF1 activity. Indeed, we demonstrate that ILC have increased IGF1R pathway activation (IGF1 ligand expression and pIGFIR levels) compared with IDC. This is similar to the results of two studies analyzing differences between ILC and IDC that found increased IGF1 ligand and IGF1R expression levels in ILC^44,45^. Consistent with this, we found that ILC cell lines are susceptible to IGF1R inhibition and importantly, that IGF1R pathway inhibitors (OSI, BMS, BEZ) synergize with a standard of care endocrine therapy (ICI) resulting in further reduced cell growth. Future studies will focus on validating these therapies in additional ILC tumors and understanding the synergistic interaction between IGF1R inhibitors and ICI. This data may be especially important given that there is an increased prevalence of late recurrences in ER+ ILC compared to ER+ IDC tumors treated with endocrine therapy, indicating the need for improved therapy options for patients with ILC^29,30^.

One limitation for the use of IGF1R inhibitors in ILC is the relatively high prevalence of mutations in the PI3K/Akt signaling pathway. Recently, Ciriello et al. comprehensively characterized ILC tumors compared to IDC tumors and described the mutational landscape of 127 ILC tumors^23^. They found that 48% of ILC harbor hotspot/missense mutations in *PIK3CA* and 13% have alterations in *PTEN,* similar to previously published data^25,45^. These genetic alterations likely lead to the elevated Akt signature they reported in these tumors. Our data suggest that the remaining tumors may also have high PI3K/Akt signaling activity due to aberrant IGF1R activity. But, because the alterations in *PIK3CA/PTEN* occur downstream of IGF1R, the effectiveness of IGF1R inhibition in this setting is unclear. Resistance to other upstream kinase inhibitors in tumors harboring activating alterations in *PIK3CA/PTEN* has been previously observed^47^ and therefore, it would be important to screen patients for these alterations before considering use of receptor tyrosine kinase inhibitor therapy. However, the use of a PI3K pathway inhibitor, such as BEZ235, may be mutually beneficial in targeting the *PIK3CA/PTEN* alterations and the enhanced IGF1R pathway activation observed in these tumors. Interestingly, Cantley et al. recently reported that high levels of insulin promote resistance to PI3K inhibitors in tumors with *PIK3CA* mutations^48^, and therefore there may also be a role for combinatorial IGF1R and PI3K inhibition. Future studies are warranted to investigate these relationships using additional *ex vivo* or *in vivo* screening of ILC tumors.

In summary, we present a diverse set of data indicating that the loss of E-cadherin enhances IGF1R pathway activity and sensitivity to IGF1R therapy, specifically in ILC. We show that IGF1R and E-cadherin directly interact, which leads to the sequestration and potential repression of IGF1R within the adherens junction. Overall, this study begins to shed light on a previously unrecognized mechanism of IGF1R regulation by E-cadherin and highlights a potential therapeutic strategy of exploiting IGF1R pathway activity in ILC tumors.

## ACKNOWLEDGEMENTS

Additional funding and support was provided by UPMC Hillman Cancer Center/Women’s Cancer Research Center. AVL and SO are recipients of Scientific Advisory Council awards from Susan G. Komen for the Cure, and AVL is a Hillman Foundation Fellow. The authors acknowledge Drs. Damir Vareslija and Leone Young for providing their expertise and media conditions for use in tumor explant culturing.

## Notes

The authors declare no potential conflicts of interest.

